# Information-Theoretic Functional Connectivity Characterizes Multiscale Network Reorganization in Postoperative Cognitive Decline

**DOI:** 10.64898/2026.05.27.728094

**Authors:** Salvatore Castelbuono, Emanuele Lo Gerfo, Gianvincenzo Sparacia, Luca Faes, Vincenzina Lo Re, Yuri Antonacci

## Abstract

Postoperative cognitive decline (POCD) after coronary artery bypass grafting (CABG) is increasingly conceptualized as a system-level disturbance of large-scale brain coordination rather than focal dysfunction. Here, we propose a multiscale neural engineering framework that combines static and dynamic information-theoretic connectivity with graph-theoretical analysis to characterize postoperative network vulnerability and its association with cognitive outcome. Resting-state fMRI was acquired in 14 male CABG patients at an early postoperative baseline (BL) and at 3-month follow-up (FU). Cognitive outcome at follow-up was assessed with the Repeatable Battery for the Assessment of Neuropsychological Status (RBANS), classifying 7 patients as POCD (RBANS < 80) and 7 as NO POCD. Functional connectivity between 32 brain regions, grouped in 8 resting-state networks (RSN), was estimated using mutual information (MI; static dependence) and mutual information rate (MIR; dynamic information exchange), each computed with parametric Gaussian (linear) and model-free *k*-nearest neighbor estimators. Pairwise connections were validated via surrogate testing, and group differences in longitudinal connectivity change (Δ = FU–BL) were assessed with permutation tests at global, intra- and inter-RSN scales. Graph metrics were computed on statistically thresholded weighted networks and related to RBANS using permutation-based Spearman correlations. POCD was not associated with a uniform reduction in connectivity but with a structured pattern of network reorganization. Static connectivity showed widespread alterations, particularly within higher-order associative systems, including salience, dorsal attention, and default mode networks. Dynamic connectivity did not exhibit global group differences but revealed selective, network-specific alterations in temporal information exchange. Longitudinal analyses showed that better cognitive outcomes were associated with increased global efficiency and density and reduced modularity and small-worldness, indicating a greater brain integration. In contrast, poorer outcomes were associated with increased segregation and higher betweenness centrality, suggesting greater reliance on hub-mediated communication. Linear measures captured more widespread connectivity changes, whereas nonlinear estimators revealed more selective alterations in dynamic information flow. Combining static and dynamic information measures with complementary estimators and surrogate-validated graph analysis reveals dissociable signatures of postoperative network dysfunction. POCD is characterized by impaired restoration of distributed integration and a progressive shift toward hub-dependent communication, suggesting that large-scale integrative vulnerability may constitute a candidate biomarker of cognitive resilience after cardiac surgery.

## 1 Introduction

Understanding how large-scale brain networks reorganize under physiological stress is a central challenge in neural engineering. Advances in functional neuroimaging and signal processing have enabled the representation of the brain as a complex network of interacting stochastic processes, paving the way for the quantitative characterization of integration, segregation and information flow across distributed systems [1, 2]. Within this context, information-theoretic measures provide a principled approach to quantifying statistical dependencies without making assumptions about the underlying distribution of the data, thereby moving beyond classical linear correlation analysis traditionally used to study brain connectivity [3]. Indeed, most clinical applications of functional connectivity (FC) still rely on static, correlation-based estimators, which quantify memoryless interactions and therefore fail to reflect properly the temporal dynamics inherent in brain data [4, 5].

Considering the information-theoretic framework, mutual information (MI) quantifies instantaneous statistical coupling between neural signals, whereas mutual information rate (MIR) extends the static measure by incorporating temporal structure and quantifying information exchange per unit time among different brain regions [3, 6]. Importantly, both measures can be estimated either using parametric formulations under the assumption that brain signals are realizations of Gaussian stochastic processes [7] or via model-free *k*-nearest neighbor (knn) estimators capable of capturing both linear and nonlinear interactions [8].

Graph-theoretical approaches are widely used to derive markers of brain information processing, even though valid inferences based on these indices critically depend on the proper assessment of the estimated connectivity patterns, regardless of the specific connectivity metric employed [9]. Moreover, the intrinsic variability of brain dynamics, particularly under pathological conditions, necessitates rigorous statistical testing to evaluate the significance of the estimated information measures. Without such validation, the reliability of these measures remains uncertain [10].

Among the various clinical conditions affecting brain function, coronary artery bypass grafting (CABG) surgery represents a clinically relevant model of systemic physiological stress, with potential consequences for large-scale brain networks. A subset of patients develops postoperative cognitive decline (POCD), characterized by impairments across memory, attention, executive, and language domains [11, 12], with important implications for functional recovery and quality of life. While POCD has traditionally been studied from a neuropsychological perspective, converging evidence indicates that postoperative cognitive vulnerability may instead reflect disruption of distributed large-scale network integration rather than focal structural damage [13, 14]. In particular, higher-order associative systems, including the default mode, salience and attentional networks, are critically involved in cognitive control, adaptive behavior, and memory processes, making them especially relevant targets for investigating network-level mechanisms of postoperative dysfunction [15, 16].

Resting-state activity itself is increasingly understood as a dynamically organized process reflecting metastable interactions among large-scale networks rather than static correlation patterns [17, 18]. Accordingly, characterizing postoperative dysfunction requires approaches capable of distinguishing between static and dynamical interactions, thereby enabling meaningful comparisons across different measures. This is particularly important because information derived from multivariate estimators can be difficult to interpret and to compare across datasets [19]. Despite growing interest in network-based biomarkers of cognitive impairment, there is a clear need to jointly examine static and dynamic brain connectivity within a unified analytical framework [20] and such integration can be achieved through the use of information-theory.

A critical but often overlooked dimension of network-based analyses concerns the spatial scale at which connectivity changes are examined. The human brain exhibits a modular, hierarchically organized architecture in which functional coupling operates simultaneously at intra-network and inter-network levels [21, 22], yet most existing studies collapse brain-wide connectivity into a single aggregate measure, thereby obscuring the scale-dependent structure of large-scale network interactions [23]. Alterations at one spatial scale need not be mirrored at another, and a purely global characterization risks confounding widespread but moderate reorganization with focal but severe disruption. To address this, we evaluate group differences across three complementary spatial scales: global connectivity, encompassing all pairwise edges; intra-RSN connectivity, restricted to within-network connections; and inter-RSN connectivity, capturing information exchange between each network and the rest of the brain [24]. This hierarchical decomposition enables the dissociation of system-wide from network-specific reorganization, and allows to determine whether postoperative cognitive vulnerability preferentially manifests as a failure of local network cohesion, inter-network coordination, or both.

Building on these considerations, we propose and apply an integrated information-theoretic framework to characterize postoperative functional network reorganization following CABG. Resting-state fMRI data are analyzed using both static (MI) and dynamic (MIR) connectivity measures, enabling the separation of instantaneous statistical coupling from temporally structured information exchange. Each measure is estimated under parametric Gaussian assumptions and via model-free knn-based approaches, allowing linear and nonlinear components of functional coupling to be disentangled, a distinction that remains actively debated in the literature [25]. All connectivity matrices are statistically validated through surrogate data analysis to ensure that retained connections reflect genuine inter-regional dependence, and are subsequently characterized using graph-theoretical metrics at both whole-brain and RSN levels. By relating longitudinal changes in network organization to neuropsychological outcomes at three months, we sought to determine whether POCD in this cohort of patients may be associated with distinct signatures of impaired static integration, altered dynamic information exchange, and large-scale topological reorganization.

## 2 Methods

### 2.1 Patients Description

Fourteen male patients (mean age 63.36 ± 9.85 years) with critical coronary artery disease scheduled for elective or urgent CABG were enrolled between February 1, 2019, and July 31, 2020. Eligible participants had been hospitalized for fewer than 6 days prior to enrollment. Patients were recruited during their stay in the intensive care unit or after transfer to the cardiothoracic unit at the *Istituto Mediterraneo per i Trapianti e Terapie ad Alta Specializzazione* (ISMETT) in Palermo, as soon as clinically feasible following surgery and after providing written informed consent. The study protocol was approved by the IRCCS ISMETT’s Institutional Research Review Board (approval number: 32/17) and the local Ethics Committee. Exclusion criteria: neurological disorders affecting lower extremity function or cognition; blindness; deafness; illiteracy; and prisoner status. All CABG procedures were performed under general anesthesia. Cardiopulmonary bypass was established via cannulation of the ascending aorta and right atrium following full systemic heparinization. Myocardial protection was achieved with warm blood cardioplegia administered every 15 minutes during aortic cross-clamp. Preoperative cognitive and motor impairments were screened through review of medical records and, when available, interviews with family members. To exclude pre-existing cognitive impairment, the Brief Intelligence Test (TIB) was administered as an estimate of premorbid intellectual functioning [26]. This test requires participants to read aloud a list of regular and irregular Italian words; since the ability to decode irregular words remains preserved following brain injury, provided that language functions are intact, the TIB yields a reliable retrospective estimate of cognitive functioning prior to the onset of illness or injury. At the three-month postoperative follow-up, cognitive performance was assessed using the Repeatable Battery for the Assessment of Neuropsychological Status (RBANS), a validated neuropsychological instrument evaluating immediate and delayed memory, attention, language, and visuospatial/constructional abilities [27]. Raw scores were converted into standardized index scores, where the normative range is defined by a mean of 100 ± 15 (1 standard deviation), with total index scores below 80 considered indicative of pathological performance, i.e. POCD.

### 2.2 fMRI Data Acquisition and Preprocessing

Functional images were acquired using a 3T MRI scanner (Discovery 750w, General Electric Medical System) equipped with a 32-channel head coil. Participants were positioned supine in the scanner, with the head stabilized using foam padding to minimize motion artifacts. Earplugs were provided to attenuate scanner noise. Resting-state fMRIs were collected using a blood oxygenation level-dependent (BOLD) echo-planar imaging sequence to measure spontaneous neuronal activity. Each resting-state session comprised 210 volumes, providing adequate temporal sampling for FC analyses. During resting-state acquisition, participants were instructed to keep their eyes closed, remain as still as possible, stay awake, and avoid engaging in structured, repetitive, or goal-directed mental activity. Baseline (BL) scans were acquired at the earliest feasible time following enrollment, predominantly within 2 days (mean 2.13 ± 2.68 days). Follow-up (FU) scans were acquired three months after surgery.

Preprocessing was performed using the CONN toolbox [28]. High-resolution T1-weighted anatomical images were used as references for spatial normalization to standard space. To ensure signal stabilization, the first 10 volumes of each functional time series were discarded. Functional images were realigned and unwarped to correct for head motion, followed by slice-timing correction to temporally align slice acquisition. Outlier volumes were identified based on frame-wise displacement (> 0.5 mm) or significant deviations in global signal intensity and were included as nuisance regressors. Physiological and motion-related confounds were removed using ordinary least squares regression applied at the voxel level, including signals from white matter and cerebrospinal fluid, six head motion parameters and their first-order temporal derivatives, and identified outlier volumes [29]. The residual BOLD time series were band-pass filtered (0.008–0.09 Hz) to isolate low-frequency fluctuations associated with resting-state connectivity while attenuating physiological and motion-related noise [30]. Spatial smoothing was performed using an 8 mm full-width at half-maximum Gaussian kernel to enhance signal-to-noise ratio and reduce inter-subject anatomical variability.

Brain parcellation was conducted in Montreal Neurological Institute (MNI) space using a predefined atlas comprising 32 regions of interest (ROIs), as defined in Supplementary Table S1. Mean BOLD time series of 200 time points were extracted from each ROI for FC analysis. The ROIs were grouped into eight canonical RSNs: Default Mode (DM; 4 ROIs), Sensorimotor (SM; 3 ROIs), Visual (VS; 4 ROIs), Salience (SAL; 7 ROIs), Dorsal Attention (DA; 4 ROIs), Frontoparietal (FP; 4 ROIs), Language (L; 4 ROIs), and Cerebellar (CB; 2 ROIs).

### 2.3 Functional Connectivity

Brain functional networks were represented as a set of *Q* = 32 stationary and ergodic stochastic processes, *S* = *{S*_1_, …, *S*_*Q*_*}*, where individual realizations of each process correspond to the BOLD time series extracted from a specific ROI (processes are assumed to have zero mean). Bivariate interactions between any pair of processes can be investigated using either static or dynamic analyses, depending on whether temporal correlations are disregarded or explicitly taken into account. Assuming [*S*_*i*_, *S*_*j*_] (*i, j* = 1, …, *Q, i ≠ j*) is a Markov process of order *m, S*_*i,n*_ and *S*_*j,n*_ denote the present states of the processes, while 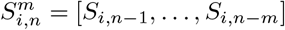 and 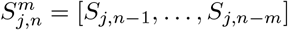 their past states.

Pairwise statistical dependencies between *S*_*i*_ and *S*_*j*_ can be investigated by means of the measure of MI, which quantifies the information shared between the variables sampling the present state of the two processes based on the concept of Shannon entropy [31]:

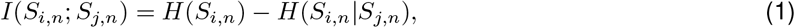

where *I*(*·*;*·*) is the MI between two random variables, *H*(*·*) denotes entropy, while *H*(*·*|*·*) represents conditional entropy.

The MIR quantifies the information exchanged per unit of time between two stochastic processes and can be regarded as a dynamic extension of MI, as it explicitly accounts for the temporal structure of the underlying interactions [32]:

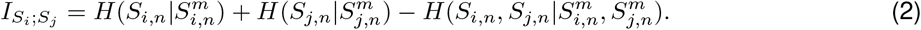

#### 2.3.1 Linear Estimator

Assuming that any pair of random variables extracted from a given bivariate process is Gaussian distributed, the analysis can be performed by exploiting linear parametric regression models. Specifically, the zero-lag dependence between any generic *S*_*i*_ and *S*_*j*_ can be expressed through the linear regression model [7]:

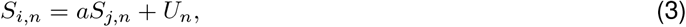

where *a* is the regression coefficient relating the two random variables and *U*_*n*_ is the residual term. Thus, the MI can be computed from the linear regression model in (3), exploiting the relationship between entropy and variance that holds for Gaussian variables [33]. In this case, the entropy of *S*_*i,n*_ can be expressed as 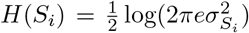, while the conditional entropy of *S*_*i*_ given *S*_*j*_ is given by 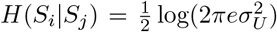, where 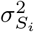 is the variance of *S*_*i*_ and 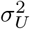 is the variance of the residual term *U*_*n*_. Substituting into (1) yields [34]:

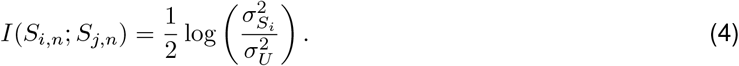

Let us now consider the generic bivariate stochastic process **S**_*n*_ = [*S*_*i,n*_, *S*_*j,n*_]^⊤^ described by the following Vector Autoregressive (VAR) Model [35]:

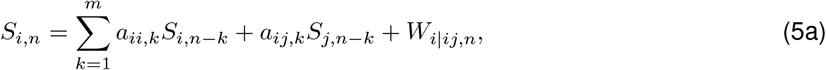

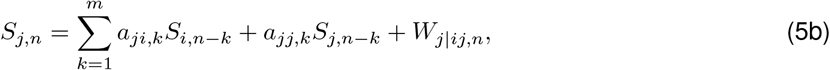

where the coefficients *a* quantify the time-lagged interactions within and between the two processes, and *W*_*i*|*ij,n*_ and *W*_*j*|*ij,n*_ are uncorrelated white noise processes with variance 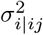 and 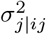. The model (5) is composed of two auto- and cross-regressive models whereby each process is regressed both on its own past and on the past of the other process. In compact form, it can be formulated as 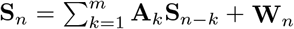, where **A**_*k*_ is a 2 *×* 2 coefficient matrix containing *a*_*ii,k*_ and *a*_*ij,k*_ in the first row and *a*_*ji,k*_ and *a*_*jj,k*_ in the second row, and **W**_*n*_ = [*W*_*i*|*ij,n*_, *W*_*j*|*ij,n*_]^⊤^ with covariance matrix 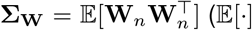, expected value). Then, the MIR between *S*_*i*_ and *S*_*j*_ can be written as follows [3]:

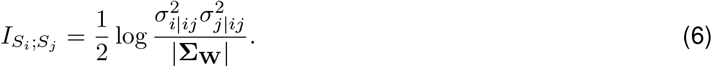

#### 2.3.2 Model-Free Estimation

The model-free estimators of MI and MIR can be computed using the *k*-nearest neighbors (knn) approach based on the KSG estimator [36]. In this framework, MI can be estimated by rewriting (1) as follows:

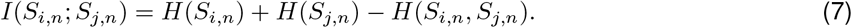

MI is obtained by searching for neighbors in the two-dimensional space [*S*_*i,n*_, *S*_*j,n*_] and performing range searches in its uni-dimensional projections. The KSG formulations facilitate the estimation of all entropy terms in (7) as follows:

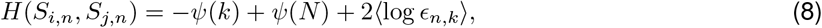

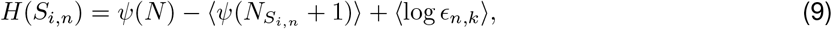

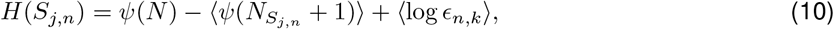

where *ψ*(*·*) is the digamma function, *N* is the total number of available bi-dimensional samples of [*S*_*i,n*_, *S*_*j,n*_], *ϵ*_*n,k*_ is twice the distance from the realizations of [*S*_*i,n*_, *S*_*j,n*_] to its *k*^*th*^ nearest neighbor, and 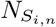 and 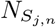 are the number of points whose distance from the realizations of *S*_*i,n*_ and *S*_*j,n*_ is strictly less than *ϵ*_*n,k*_*/*2.

Substituting the terms obtained in (8), (9) and (10) into (7) an estimate of the MI is obtained as follows:

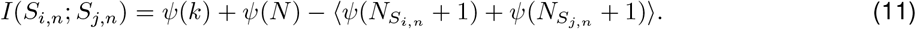

The estimation of MIR requires instead six entropy terms as each conditional entropy term in (2) is the difference between two entropies [37]. These terms can be estimated as follows:

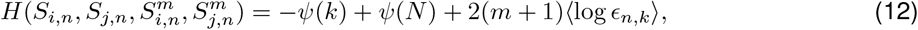

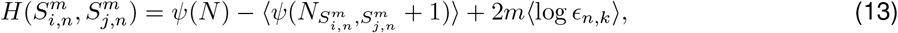

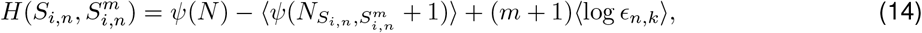

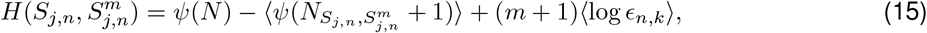

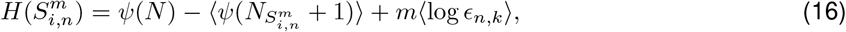

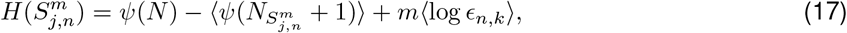

where *N* is the total number of available 2(*m* + 1)-dimensional points 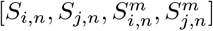, and 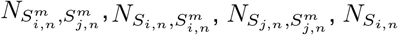 and 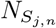 are the number of points whose distance from the correspondent reference points is smaller than *ϵ*_*n,k*_*/*2 in the lower dimensional spaces

(i.e., 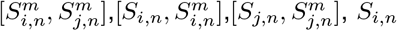, and 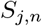 respectively), being *ϵ*_*n,k*_ twice the distance from samples in 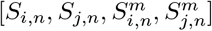 to the *k*-th neighbors. Thus, an estimate of the MIR can be obtained as:

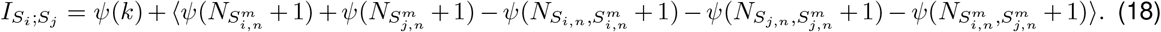

This nonparametric formulation enables estimation of entropy rates without Gaussian assumptions, allowing detection of nonlinear and time-dependent interactions between ROIs. For the purpose of this work, a value of *k* = 10 nearest neighbors [38] was chosen. To account for potential bias, the results were computed using different values of *m* ∈ {1, 3, 5}.

#### 2.3.3 Statistical Analysis

To assess whether longitudinal FC changes differed between patients who developed POCD and those who did not (NO POCD), the difference matrix for each participant was first computed as the difference between Follow Up and Baseline (Δ*FC* = FU − BL). Group differences were evaluated at three spatial scales: global connectivity, i.e. considering all edges; intra-RSN connectivity, considering only within-network edges; and inter-RSN connectivity, considering edges between a RSN and all the others. For each scale and connectivity estimator, the median Δ connectivity was computed separately for POCD and NO POCD groups. Statistical significance of between-group differences was assessed using a nonparametric permutation test, with number of permutation equal to 5000. Specifically, group labels (POCD vs NO POCD) were randomly permuted across subjects to generate the empirical null distribution of the difference in median Δ under the null hypothesis of no group effect. Two-tailed *p*-values were computed as the proportion of permuted statistics exceeding the observed difference in absolute magnitude. Statistical significance was set at *α* = 0.05.

### 2.4 Surrogate Data Analysis

The statistical significance of each estimate of pairwise connectivity was performed using a surrogate data approach testing the null hypothesis of no inter-regional coupling between *S*_*i*_ and *S*_*j*_ [39]. For each ROI pair, *N*_*s*_ = 500 surrogate realizations were generated using a circular shift approach. This procedure preserves the internal dynamical structure of each time series while destroying any cross-dependence between them. For each original pair (*S*_*i*_, *S*_*j*_), both MI and MIR were computed using the linear Gaussian formulation and the model-free *k*-nearest neighbor estimator. The same estimators were then exploited for each surrogate pair to generate empirical null distributions under the hypothesis of independent processes with preserved univariate dynamics. For each connectivity metric (MI linear, MI knn, MIR linear, MIR knn), surrogate-based null distributions were obtained from the *N*_*s*_ realizations. Significance thresholds were defined as the 100(1 − *α*)^*th*^ percentile of the surrogate distribution with *α* = 0.01. An observed connectivity value was deemed statistically significant if it exceeded the corresponding threshold. This procedure was repeated independently for each ROI pair and each connectivity estimator. The surrogate-based inference framework ensures that detected connections reflect genuine inter-regional information exchange beyond what can be explained by individual spectral properties or autocorrelation structure alone.

### 2.5 Graph-Theoretical Analysis

For each participant (*N* = 14), connectivity matrices derived from linear MI, knn-based MI, linear MIR, and knn-based MIR were computed at baseline and follow-up, and statistically thresholded against their surrogate-derived significance matrices, yielding weighted undirected adjacency matrices retaining only significant connections. Graph-theoretical measures were then computed on the resulting functional brain networks (*Q* = 32 nodes) [40]. The network is modeled as a graph *G* = (*V, E*), with weighted adjacency matrix *W* = [*w*_*ij*_] and binary support *A* = [*a*_*ij*_], where *a*_*ij*_ = 1 if *w*_*ij*_ > 0, *a*_*ii*_ = *w*_*ii*_ = 0 otherwise. For path-based measures, edge weights were mapped to lengths ℓ_*ij*_ = 1*/w*_*ij*_ (ℓ_*ij*_ = *∞* if *w*_*ij*_ = 0), so that stronger connections correspond to shorter topological distances, and shortest path distances *d*_*ij*_ were computed on *L* = [ℓ_*ij*_]. The following measures were extracted for each adjacency matrix.

*Mean nodal degree* (K) and *mean nodal strength* (S) index overall network connectedness and interaction intensity, respectively:

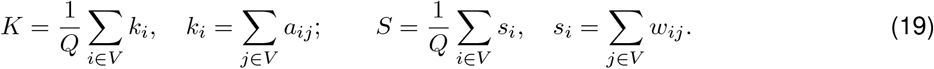

*Density* is the fraction of existing edges *N*_*E*_ over all possible undirected edges:

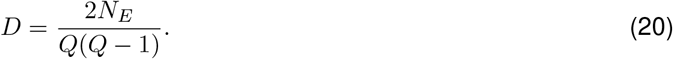

*Mean betweenness centrality* reflects the average reliance on intermediary nodes for shortest-path communication:

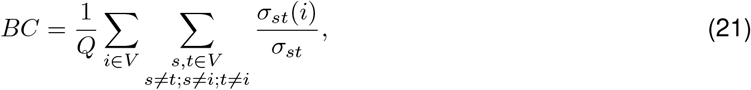

where *σ*_*st*_ is the number of shortest weighted paths between *s* and *t*, and *σ*_*st*_(*i*) the subset passing through *i*.

*Global* and *local efficiency* quantify integration and segregation of information transfer across the network:

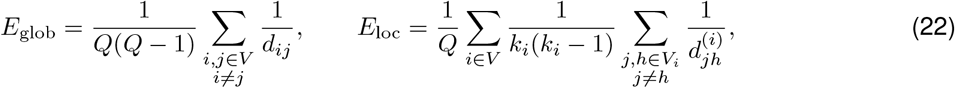

where *V*_*i*_ is the neighborhood of node *i* and 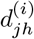 the shortest-path distance between *j* and *h* within that subgraph. Disconnected pairs contribute zero to both sums (1*/* ∞ = 0).

*Modularity* quantifies the degree to which the network segregates into densely intra-connected commu-nities:

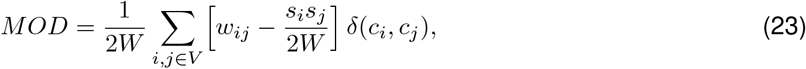

Where 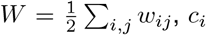 is the community assignment of node *i*, and *δ*(·, ·) is the Kronecker delta (equal to 1 if *c*_*i*_ = *c*_*j*_, 0 otherwise).

*Small-world index* characterizes whether the network exhibits small-world organization, i.e. higher clustering and comparable path lengths relative to a random topology of equivalent size and degree sequence:

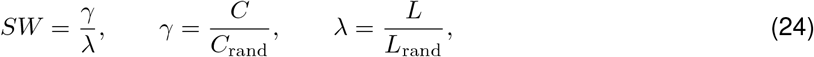

where *C* is the mean weighted clustering coefficient,

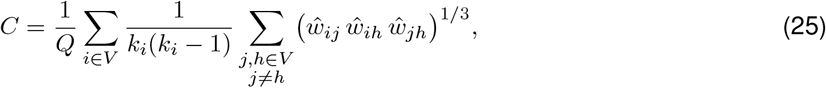

with *ŵ*_*ij*_ = *w*_*ij*_*/* max(*w*), and *L* the characteristic path length averaged over reachable node pairs only:

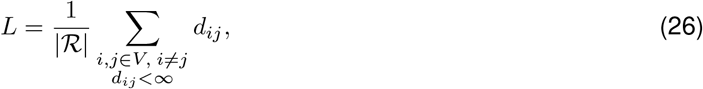

where |ℛ|= |{(*i, j*) : *i* ≠ *j, d*_*ij*_ <∞}| is the number of reachable ordered node pairs. Reference values *C*_rand_ and *L*_rand_ were averaged over *N*_rand_ = 100 surrogate networks generated by degree-preserving random rewiring, applying the same exclusion criterion for disconnected pairs when estimating *L*_rand_. *γ* > 1 indicates above-random clustering, *λ*≈ 1 indicates path lengths comparable to a random network and *SW* > 1 is the hallmark of small-world organization.

For each measure, a delta score (Δ = FU −BL) was computed per participant to capture longitudinal change.

### 2.6 Association Between Network Reorganization and Cognitive Outcome

To investigate the relationship between longitudinal network reorganization and cognitive outcome, Spearman’s rank correlation coefficients *ρ* were computed between changes in graph-theoretic metrics (Δ = FU − BL) and RBANS total score for each connectivity estimator (MI linear, MI knn, MIR linear, MIR knn). Given the limited sample size (*N* = 14), statistical significance of Spearman’s *ρ* was assessed using a nonparametric permutation test. Specifically, RBANS scores were randomly permuted (*N*_*p*_ = 5000) across subjects to generate the empirical null distribution of *ρ*, and two-tailed *p*-values were computed as the proportion of permuted coefficients exceeding the observed value in absolute magnitude. Statistical significance was set at *α* = 0.05. To further assess robustness and sampling variability, a subject-level bootstrap analysis (5000 resamples with replacement) was performed. For each bootstrap sample, correlations between RBANS scores and graph metrics were recomputed, and percentile-based 95% confidence intervals were derived. In addition, a sign-stability index was calculated as the proportion of bootstrap samples in which the correlation coefficient retained the same sign as the original estimate. This complementary resampling strategy allowed us to evaluate both statistical significance and robustness of the observed associations under small-sample conditions.

## 3 Results

### 3.1 Participant Characteristics

At follow-up, 7 patients met criteria for POCD (RBANS 73.71 ± 5.02), while 7 patients were classified as NO POCD (RBANS 98.14 ± 12.29). No significant differences were observed between POCD and NO POCD patients in age (62.72 ± 11.18 vs 64.00 ± 9.18 years; *p* = 0.418). As expected, the groups differed significantly in RBANS total score at follow-up (*p* < 0.001), confirming the validity of the classification criterion.

### 3.2 Static and Dynamic Connectivity Alterations at Multiscale Level

Figure 1 illustrates the median longitudinal changes in functional connectivity for the POCD and NO POCD groups obtained using both the linear and model-free MI estimators. Results of the permutation analyses are summarized in Table 1. At the global level, statistically significant between-group differences are observed for both the parametric linear estimator and the knn-based nonparametric estimator. Specifically, for the linear MI measure, the NO POCD group exhibits 318 positive and 178 negative edges, whereas the POCD group showed 206 positive and 290 negative edges. Comparable patterns emerge for the knn-based MI, with 295 positive and 201 negative edges in the NO POCD group versus 208 positive and 288 negative edges in the POCD group. At the network level, the salience network displays high significant (*p* < 0.01) between-group differences for both estimators. Additionally, the linear estimator identifies statistically significant alterations within the dorsal attention network (*p* < 0.05), while the model-free estimator detects significant effects within the frontoparietal network (*p* < 0.05). At the inter-network scale, both methods consistently indicate that connectivity patterns involving the default mode network are the most strongly affected (*p* < 0.01).

**Table 1:** Results of permutation testing comparing longitudinal functional connectivity changes (Δ*FC* = FU− BL) between POCD and NO POCD groups for Mutual Information (MI) and Mutual Information Rate (MIR; *m* = 3), using linear and knn-based estimators, at global, intra-RSN, and inter-RSN scales. Statistical significance is indicated by asterisks (*∗*: *p* < 0.05; *∗∗*: *p* < 0.01).

|  | MI linear | MI knn | MIR linear | MIR knn |
| --- | --- | --- | --- | --- |
| Global FC | **0.011\*** | **0.004\*\*** | 0.156 | 0.258 |
| Intra-RSN |  |  |  |  |
| DM | 0.178 | 0.315 | 0.635 | 0.251 |
| SM | 0.620 | 0.499 | 0.495 | 0.129 |
| VS | 0.323 | 0.104 | 0.478 | 0.264 |
| SAL | **0.007\*\*** | **0.006\*\*** | **0.048\*** | 0.689 |
| DA | **0.021\*** | 0.796 | **0.005\*\*** | 0.298 |
| FP | 0.305 | **0.016\*** | 0.766 | 0.362 |
| L | 0.699 | 0.305 | 0.704 | 0.344 |
| CB | 0.609 | 0.772 | 0.725 | 0.884 |
| Inter-RSN |  |  |  |  |
| DM | **0.009\*\*** | **0.001\*\*** | 0.332 | 0.177 |
| SM | 0.254 | 0.781 | 0.841 | 0.773 |
| VS | 0.369 | 0.751 | 0.195 | 0.381 |
| SAL | 0.421 | 0.494 | 0.498 | 0.255 |
| DA | 0.329 | 0.447 | 0.247 | **0.049\*** |
| FP | 0.125 | 0.199 | 0.514 | 0.896 |
| L | 0.369 | 0.202 | 0.488 | 0.315 |
| CB | 0.230 | 0.079 | 0.314 | 0.203 |

**Figure 1:**
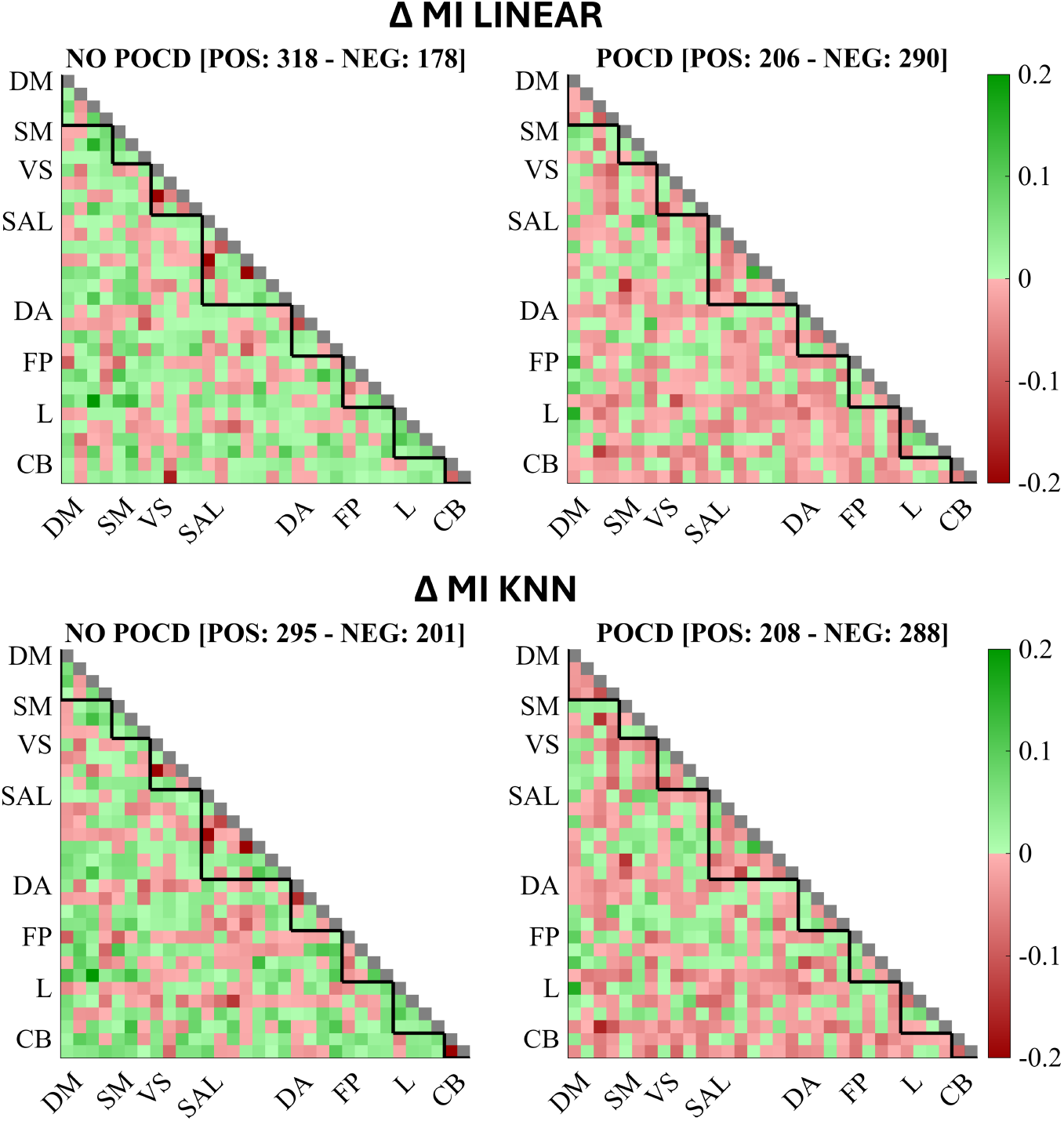
Median Δ FC matrices for the two groups NO POCD (left column) and POCD (right column) estimated as linear (top row) and knn-based model-free (bottom row) MI. Edges with a positive difference are reported in green, while edges with negative changes in red. Number of positive (POS) and negative (NEG) edges are reported in the figure. Black triangles show intra-RSN FC portion.

Results for MIR are here presented for *m* = 3; analyses using *m* ∈ {1, 5} are reported in Supplementary Table S2 and Figures S1–S2. Figure 2 depicts the median longitudinal changes in functional connectivity for the POCD and NO POCD groups estimated using both linear and model-free MIR approaches. The corresponding permutation test results are reported in Table 1. At the global level, no significant between-group differences are detected for either the parametric linear estimator or the knn-based nonparametric estimator. For the linear MIR metric, both groups show a predominance of edges with negative median Δ*FC*, with a more pronounced shift toward negative values in the POCD group. Specifically, the NO POCD group exhibits 217 positive and 279 negative edges, whereas the POCD group shows 168 positive and 328 negative edges. A similar pattern is observed for the knn-based MIR estimator, although the longitudinal changes are smaller in absolute magnitude compared with both MI and linear MIR. In this case, the NO POCD group presents 271 positive and 225 negative edges, while the POCD group shows 185 positive and 311 negative edges. At the network level, only the linear MIR estimator with *m* = 3 reveals a significant between-group differences, localized within the dorsal attention network (*p* < 0.01) and salience network (*p* < 0.05), whereas the knn-based MIR estimator does not identify any significant intra-network effects for each value of *m* considered. At the inter-network level, linear MIR does not detect significant connectivity differences, while the MIR knn estimated with *m* = 3 indicates effects involving DA-related connections (*p* < 0.05). Across all estimators, the NO POCD group consistently displays a higher proportion of edges with positive longitudinal change (Δ*FC* > 0), whereas the POCD group exhibits a greater number of edges with negative Δ*FC* (Figures 1 and 2).

**Figure 2:**
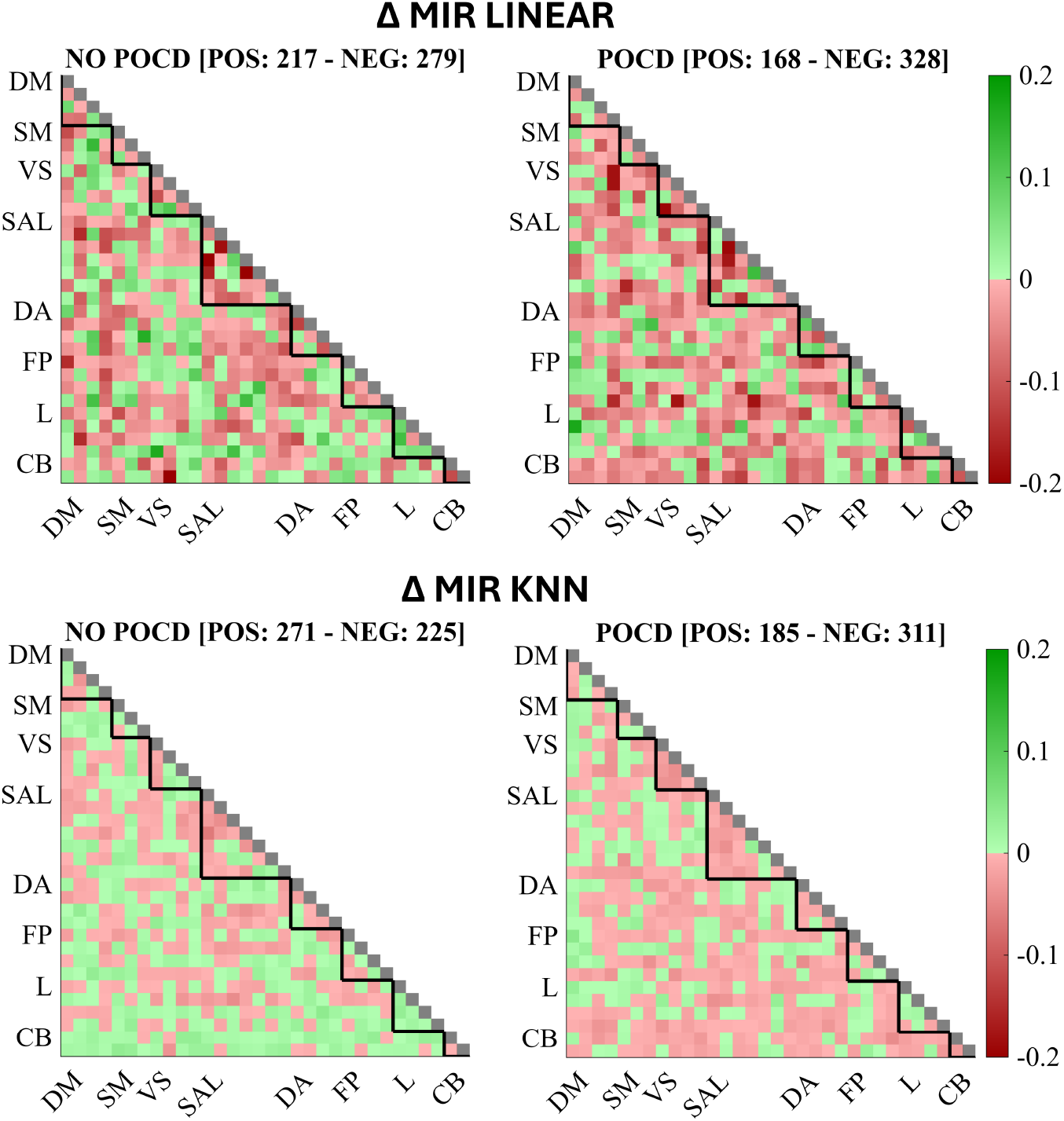
Median Δ FC matrices for the two groups NO POCD (left column) and POCD (right column) estimated as linear (top row) and knn-based model-free (bottom row) MIR with *m* = 3. Edges with a positive difference are reported in green, while edges with negative changes in red. Number of positive (POS) and negative (NEG) edges are reported in the figure. Black triangles show intra-RSN FC portion.

### 3.3 Longitudinal Network Reorganization and Cognitive Outcome

Surrogate-thresholded FC matrices are constructed at baseline and follow-up for each participant across all four estimators; the number of significant pairwise connections for each estimator is reported in Supplementary Table S3. Graph-theoretical metrics derived from these networks show systematic associations between longitudinal reorganization and cognitive outcome at three months. Only statistically significant associations are reported here; complete results for all metrics and estimators are provided in Supplementary Table S4. Figure 3 reports statistically significant correlations for the linear MI estimator, which yields the largest number of significant associations: RBANS scores are positively correlated with changes in network density (Fig. 3a; *ρ* = 0.56, *p* < 0.05) and global efficiency (Fig. 3c; *ρ* = 0.66, *p* < 0.05), and negatively correlated with modularity (Fig. 3b; *ρ* = −0.72, *p* < 0.05) and small-world index (Fig. 3d; *ρ* = −0.62, *p* < 0.05). The knn-based MIR estimator additionally reveals significant negative correlations between RBANS scores and changes in mean betweenness centrality (Fig. 4a; *ρ* = −0.65, *p* < 0.05) and small-world index (Fig.4b; *ρ* = −0.58, *p* < 0.05). Bootstrap resampling supports the robustness of these associations. For linear MI, the positive relationships with Δ density (95% CI [0.001, 0.897], stability = 0.97) and Δ global efficiency (95% CI [0.123, 0.881], stability = 0.99) are stable, as are the negative associations with Δ modularity (95% CI [−0.924, −0.227], stability = 1.00) and Δ small-world index (95% CI [−0.886, −0.036], stability = 0.98), all showing high sign stability (≥0.97). Similarly, knn-based MIR yields robust negative associations for Δ betweenness centrality (95% CI [−0.893, −0.205], stability = 0.99) and Δ small-world index (95% CI [−0.954, −0.049], stability = 0.98). In contrast, no associations derived from knn-based MI or linear MIR retain confidence intervals excluding zero, suggesting weaker or less stable effects for these estimators.

**Figure 3:**
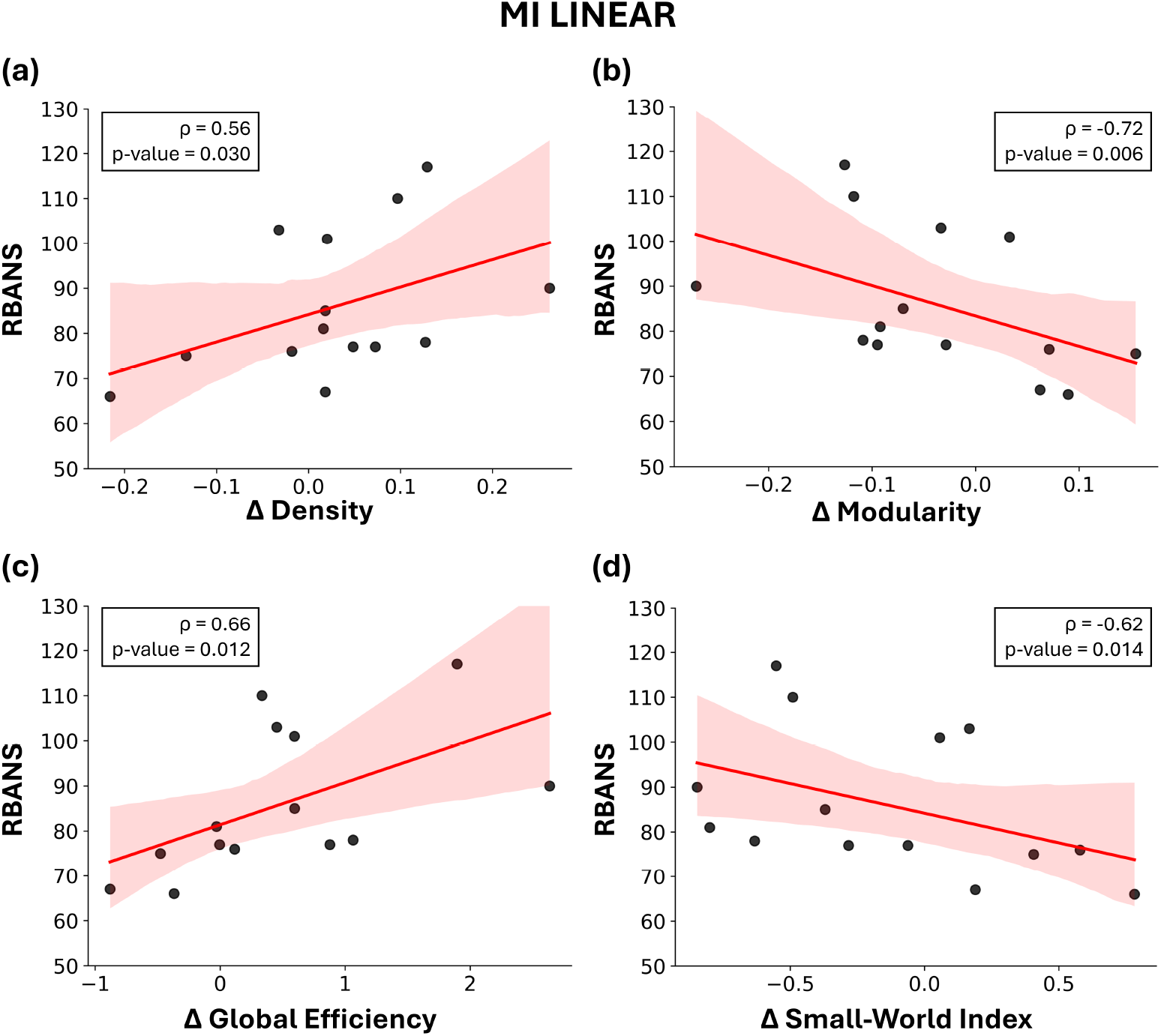
Significant associations between RBANS scores and longitudinal changes (Δ = FU − BL) in graph-theoretical metrics derived from functional connectivity based on linear Mutual Information (MI). Panels show: (a) Δ density, (b) Δ modularity, (c) Δ global efficiency, and (d) Δ small-world index. Red shaded areas represent the 95% confidence intervals. Insets report Spearman’s correlation coefficients (*ρ*) and corresponding *p*-values.

**Figure 4:**
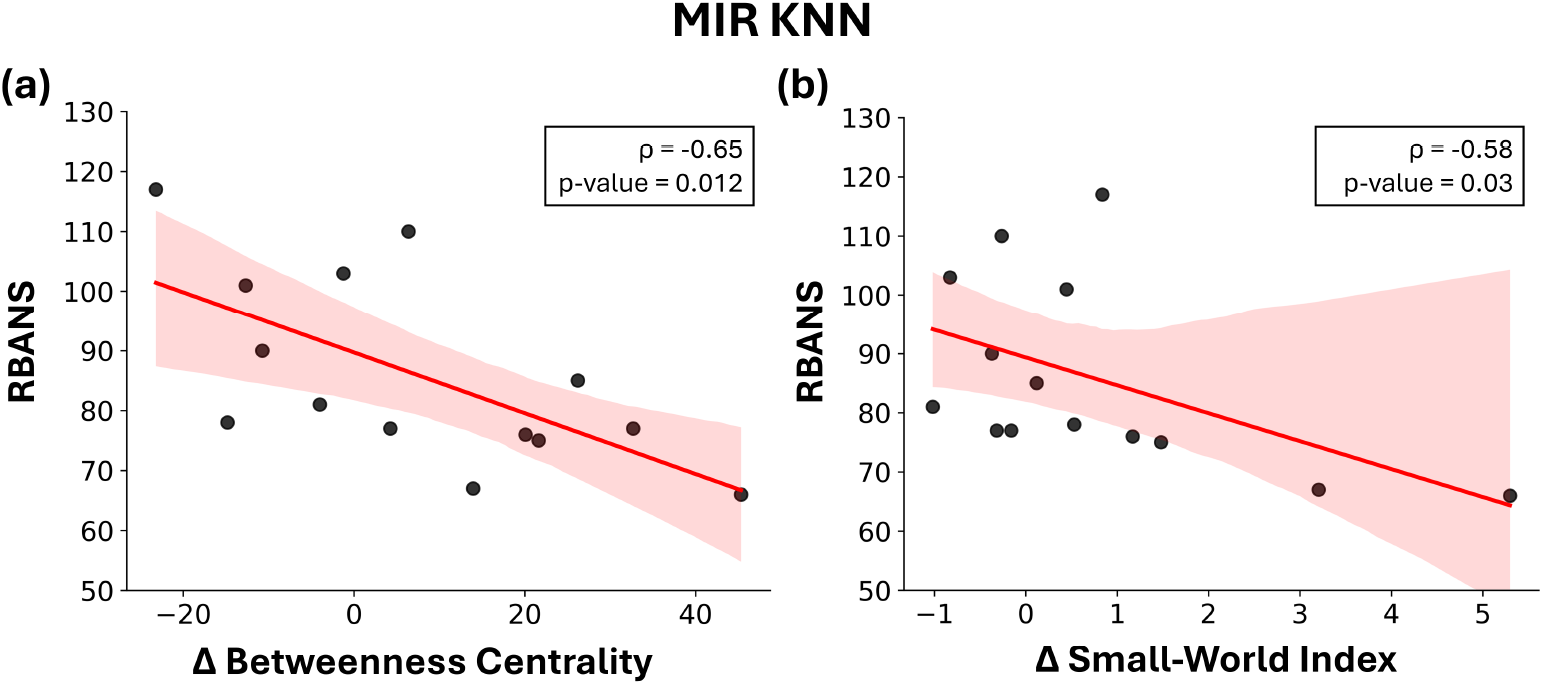
Significant associations between RBANS scores and longitudinal changes (Δ = FU−BL) in graph-theoretical metrics derived from functional connectivity based on knn-based Mutual Information Rate (MIR). Panels show: (a) Δ betweenness centrality and (b) Δ small-world index. Red shaded areas represent the 95% confidence intervals. Insets report Spearman’s correlation coefficients (*ρ*) and corresponding *p*-values.

## 4 Discussion

The present study investigated postoperative functional network reorganization in patients undergoing CABG and its relationship with cognitive outcome at three months. By combining static and dynamic information-theoretic connectivity measures with graph-theoretical analysis across global and RSN scales, we aimed to characterize the multilevel structure of postoperative brain reconfiguration. Our results suggest that POCD is not associated with a uniform reduction in connectivity, but rather with a structured pattern of altered integration, segregation and hub organization. Differences between POCD and NO POCD patients emerged across static and dynamic domains, across linear and model-free estimators, and across specific large-scale associative systems. Collectively, these findings suggest that POCD reflects a systems-level disturbance of large-scale brain coordination rather than isolated regional dysfunction.

### 4.1 Patterns in POCD Functional Connectivity Dysfunction

At the static level, POCD patients exhibited significant global and RSN-level differences compared to NO POCD, with a predominance of negative longitudinal edge changes. These effects were most prominent within and between higher-order associative systems, particularly the salience, dorsal attention, and default mode networks. Such findings may indicate an impaired restoration of large-scale statistical interdependence following surgery, a phenomenon increasingly recognized as a hallmark of postoperative neurocognitive impairment [41]. Convergent evidence from aging and prodromal neurodegenerative populations similarly links reduced inter-network coupling among associative systems to heightened cognitive vulnerability and reduced adaptive capacity [42, 43].

The RSNs most affected form a coordinated control architecture. The salience network detects behaviorally relevant stimuli and orchestrates switching between the internally oriented processes mediated by the default mode system and the externally directed control systems [15, 44]; disruption within SAL or in its inter-network coupling may therefore reflect impaired allocation of cognitive resources. The dorsal attention network supports sustained top-down attentional modulation [45], and altered connectivity within this system may contribute to the attentional instability frequently observed in POCD. The default mode network, a central integrative hub, underpins memory consolidation, self-referential processing and internally directed cognition [16]; perturbations in its coupling with task-positive systems suggest impaired coordination between executive and mnemonic processes, consistent with the variability in RBANS performance observed here.

Dynamic analyses using MIR revealed a distinct and complementary spatial profile. No significant global group differences were detected for either MIR estimator, indicating that the overall rate of dynamic information exchange across the network may be comparably affected in both groups following surgery. Nevertheless, edge distributions showed a higher proportion of negative ΔFC values in the POCD group, consistent with a subtle, diffuse attenuation that did not reach statistical significance at the global scale. Despite this absence of global effects, MIR identified localized alterations: linear MIR highlighted significant intra-network changes within the dorsal attention and salience networks, whereas nonlinear MIR revealed inter-network effects, particularly involving dorsal attention-related connections. Notably, this pattern was not replicated at *m* = 1 or *m* = 5, suggesting that the temporal scale of embedding governs which aspects of dynamic reorganization become detectable, as we discuss in Section 4.3.

This dissociation between static and dynamic profiles is consistent with network vulnerability models proposing that hub-centered systems, particularly those anchored in the default mode network, are disproportionately susceptible to metabolic, inflammatory, or vascular stressors [22, 46]. Because such hubs support communication across distributed subsystems, their perturbation can induce widespread reorganization in statistical coupling without globally suppressing dynamic information transfer. The emergent picture is therefore one of spatial selectivity: static connectivity is broadly disrupted across higher-order systems, whereas dynamic reorganization is circumscribed within specific network interactions, suggesting that the two domains capture partially independent dimensions of postoperative network dysfunction.

### 4.2 The Global Remodulation of the Functional Network in POCD

Longitudinal changes in graph-theoretical properties provided a complementary, topology-level characterization of the network alterations described above. When connectivity was estimated using linear MI, RBANS scores were positively associated with changes in network density and global efficiency, and negatively associated with changes in modularity and small-world index; knn-based MIR additionally revealed negative associations between RBANS and changes in betweenness centrality and small-world index.

Taken together, these associations delineate two contrasting trajectories of network reconfiguration. In patients with better cognitive outcomes, the network shifted toward higher density and global efficiency, reflecting an expansion of distributed connectivity and more efficient long-range communication [1, 40]. Concurrently, these patients exhibited lower modularity and a reduced small-world index, indicating a partial dissolution of rigid community structure in favor of greater inter-network integration. This configuration is compatible with a partial restoration of the flexible, weakly modular topology that characterizes healthy resting-state organization and supports efficient large-scale processing [1]. In contrast, patients with poorer cognitive outcomes exhibited increased modularity alongside greater betweenness centrality. Elevated modularity reflects stronger within-module cohesion at the expense of inter-network communication, consistent with a fragmentation of large-scale systems rather than adaptive specialization [47]. Increased betweenness centrality, in turn, indicates progressive centralization of network communication through a restricted set of hub regions. While hubs are essential for efficient information routing, excessive reliance on them reduces redundancy and increases topological vulnerability to perturbation [40, 48, 49].

The coexistence of segregation and hub centralization in poorer-outcome patients therefore points to a maladaptive reconfiguration in which information processing becomes simultaneously compartmentalized and overly dependent on specific relay nodes, a pattern qualitatively distinct from the integrative restoration observed in cognitively preserved patients. This network profile resembles alterations reported in aging and early neurodegeneration. Conditions such as mild cognitive impairment and Alzheimer’s disease are characterized by reduced global efficiency, disrupted hub organization and increased topological vulnerability [42, 43, 49, 50]. The convergence with these profiles supports the interpretation of POCD as a condition of accelerated topological vulnerability, in which surgery acts as a precipitating stressor that reveals, or transiently amplifies, a pre-existing susceptibility toward less resilient network configurations, rather than constituting an isolated surgical complication.

### 4.3 Linear vs Nonlinear Functional Connectivity

A further relevant aspect concerns the comparison between parametric (linear) and model-free (knn-based) estimations for both static MI and dynamic MIR. Although both estimators quantify statistical dependence, the linear formulation relies on Gaussian assumptions and captures dependence through covariance structure, whereas the knn estimator is model-free and sensitive to any form of interaction, including linear, nonlinear, or mixed dependencies. Under Gaussian assumptions, MI reduces to a function of squared correlation coefficient, thereby reflecting second-order statistics [7, 31]. In contrast, knn-based estimator directly approximate entropy from the data distribution without parametric constraints, allowing detection of more complex coupling patterns [8].

For static connectivity, linear estimation revealed broader global and intra-network differences between POCD and NO POCD patients, suggesting that functional coupling is widely affected following surgery. The knn-based MI revealed a similar pattern of alterations, albeit with greater spatial selectivity, particularly at the inter-network level, indicating that nonlinear dependencies may reorganize in a more localized fashion. A similar, though not identical, dissociation was observed for dynamic connectivity. MIR explicitly incorporates temporal structure and reflects the rate of information exchange over time [51]. Linear MIR highlighted more extensive alterations in dynamic coupling, whereas knn-based MIR revealed more circumscribed inter-network differences among associative control systems. This suggests that postoperative cognitive decline affects both the magnitude of temporal coordination, captured by linear estimators, and the structural complexity of dynamic information transfer, captured by model-free estimation. The convergence and divergence between linear and knn estimators across both MI and MIR indicate that POCD-related reorganization operates across multiple statistical layers of functional connectivity, with linear measures appearing more sensitive to widespread alterations.

This is consistent with accumulating evidence that linear models faithfully describe macroscopic brain dynamics, in particular at the spatiotemporal scale of fMRI [25]. This effective macroscale linearity emerges from spatial averaging across large neuronal populations and temporal averaging through the hemodynamic response function, both of which suppress microscopic nonlinear dynamics. Consequently, fMRI signals are often well approximated by low-order autoregressive descriptions, a property supported by recent large-scale benchmarking studies showing that nonlinear model components confer limited predictive advantage over linear counterparts, even when explicitly optimized for resting-state data [52]. The broader sensitivity of linear MI and MIR estimators to POCD-related alterations therefore likely reflects the genuine linear character of macroscale resting-state coupling, rather than a limitation of the parametric approach.

A related methodological observation concerns the sensitivity of MIR-based results to the model order *m*. While analyses conducted at *m* = 3 revealed significant intra-network differences within the dorsal attention and salience networks, the corresponding analyses at *m* = 1 and *m* = 5 did not yield significant between-group effects at any spatial scale (Supplementary Table S2). This dependence on *m* should not be interpreted solely as a statistical instability: the model order governs the temporal horizon over which past states are incorporated into the MIR estimate, and therefore determines which timescale of dynamic information exchange is effectively characterized. The absence of significant effects at *m* = 1 may reflect an insufficient embedding depth to capture the memory structure of resting-state BOLD signals, whereas at *m* = 5 the increased dimensionality of the embedding space may reduce estimation precision given the limited time series length available (200 time points), a well-known constraint of nonparametric entropy estimators in high-dimensional settings [8].

### 4.4 Limitations

Several limitations should be considered when interpreting these findings. First, the sample size was relatively small (*N* = 14), limiting statistical power and generalizability. Moreover, the absence of a comprehensive preoperative neuropsychological baseline precludes strict inference of individual-level cognitive decline. Replication in larger cohorts with longitudinal cognitive assessment is necessary to confirm the robustness of the observed network signatures. Second, the cohort consisted exclusively of male participants, and sex-dependent biological factors, including hormonal profiles and neuroinflammatory responses, can significantly modulate both baseline network topology and susceptibility to cognitive decline; future research involving sex-balanced samples is therefore essential. Third, the large number of comparisons across estimators, spatial scales, and graph metrics increases the risk of false-positive findings; no correction for multiple comparisons was applied given the exploratory nature of the study, and results should be interpreted accordingly. Finally, the absence of a non-surgical control group prevents dissociation between surgery-related network changes and natural longitudinal variability in functional connectivity, a gap that future studies should address by incorporating matched controls.

## 5 Conclusion

The present findings suggest that postoperative cognitive decline following CABG may be associated with a structured reorganization of functional brain networks, rather than a uniform reduction in connectivity. In this cohort, POCD was characterized by patterns consistent with reduced large-scale integration, increased segregation, and a tendency toward greater hub centralization, particularly within higher-order associative systems. Conversely, preserved cognitive outcomes were associated with network configurations indicative of more efficient and distributed communication. By combining static and dynamic information-theoretic measures with both linear and model-free estimators, these findings provide preliminary evidence that post-operative network alterations may span multiple statistical and temporal scales. Linear measures appeared to capture more widespread changes in functional coupling, whereas nonlinear dynamic approaches highlighted more selective alterations in the organization of information flow. Notably, the observed network patterns show similarities with those reported in conditions such as mild cognitive impairment, Alzheimer’s disease, and vascular cognitive impairment. While this convergence should be interpreted with caution, it is consistent with the hypothesis that POCD may involve a shift toward a more vulnerable large-scale network configuration. Within the framework of perioperative neurocognitive disorders, these results support a systems-level interpretation of postoperative cognitive dysfunction. Given the limited sample size and cohort characteristics, these findings should be considered exploratory. Future studies in larger and more heterogeneous populations will be necessary to confirm the robustness of these network signatures and to evaluate their potential role as biomarkers of cognitive vulnerability and recovery after surgery.

## Author Contributions

S.C.: Conceptualization, Methodology, Software, Formal analysis, Visualization, Writing - original draft. E.L.G.: Data curation, Neuropsychological assessment, Writing - review & editing. G.S.: Data curation, MRI acquisition and supervision, Writing - review & editing. L.F.: Conceptualization, Methodology, Supervision, Writing - review & editing. V.L.R.: Clinical supervision, Resources, Project administration, Writing - review & editing. Y.A.: Conceptualization, Methodology, Supervision, Writing - review & editing. V.L.R. and Y.A. contributed equally to this work.

## Data Availability

The data that support the findings of this study are not publicly available due to ethical restrictions and patient privacy regulations governing clinical research conducted at IRCCS ISMETT (approval number: 32/17). De-identified data may be made available upon reasonable request to the corresponding author, subject to approval by the local Ethics Committee.

## References

[1] Ed Bullmore and Olaf Sporns. Complex brain networks: graph theoretical analysis of structural and functional systems. Nature Reviews Neuroscience, 10(3):186–198, 2009. doi: 10.1038/nrn2575.

[2] Danielle S. Bassett and Olaf Sporns. Network neuroscience. Nature Neuroscience, 20(3):353–364, 2017. doi: 10.1038/nn.4502.

[3] Gorana Mijatovic, Yuri Antonacci, Michal Javorka, Daniele Marinazzo, Sebastiano Stramaglia, and Luca Faes. Network representation of higher-order interactions based on information dynamics. IEEE Transactions on Network Science and Engineering, 12(3):1872–1884, 2025. doi: 10.1109/TNSE.2025.3540982.

[4] Karl J. Friston. Functional and effective connectivity: a review. Brain Connectivity, 1(1):13–36, 2011. doi: 10.1089/brain.2011.0008.

[5] Luca Faes, Laura Sparacino, Gorana Mijatovic, Yuri Antonacci, Leonardo Ricci, Daniele Marinazzo, and Sebastiano Stramaglia. Partial information rate decomposition. Phys. Rev. Lett., 135:187401, Oct 2025. doi: 10.1103/nrwj-n8lj. URL https://link.aps.org/doi/10.1103/nrwj-n8lj.

[6] Luca Faes, Alberto Porta, and Giandomenico Nollo. Information decomposition in bivariate systems: Theory and application to cardiorespiratory dynamics. Entropy, 17(1):277–303, 2015. ISSN 1099-4300. doi: 10.3390/e17010277. URL https://www.mdpi.com/1099-4300/17/1/277.

[7] Luca Faes, Silvia Erla, and Giandomenico Nollo. Measuring connectivity in linear multivariate processes: Definitions, interpretation, and practical analysis. Computational and Mathematical Methods in Medicine, 2012(1):140513, 2012. doi: 10.1155/2012/140513. URL https://onlinelibrary.wiley.com/doi/abs/10.1155/2012/140513.

[8] Chiara Barà, Laura Sparacino, Riccardo Pernice, Yuri Antonacci, Alberto Porta, Dimitris Kugiumtzis, and Luca Faes. Comparison of discretization strategies for the model-free information-theoretic assessment of short-term physiological interactions. Chaos: An Interdisciplinary Journal of Nonlinear Science, 33(3):033127, 03 2023. ISSN 1054-1500. doi: 10.1063/5.0140641. URL https://doi.org/10.1063/5.0140641.

[9] Jlenia Toppi, Donatella Mattia, Monica Risetti, Rita Formisano, Fabio Babiloni, and Laura Astolfi. Testing the significance of connectivity networks: Comparison of different assessing procedures. IEEE Transactions on Biomedical Engineering, 63(12):2461–2473, 2016. doi: 10.1109/TBME.2016.2621668.

[10] Helder Pinto, Ivan Lazic, Yuri Antonacci, Riccardo Pernice, Danlei Gu, Chiara Barà, Luca Faes, and Ana Paula Rocha. Testing dynamic correlations and nonlinearity in bivariate time series through information measures and surrogate data analysis. Frontiers in Network Physiology, Volume 4-2024, 2024. ISSN 2674-0109. doi: 10.3389/fnetp.2024.1385421. URL https://www.frontiersin.org/journals/network-physiology/articles/10.3389/fnetp.2024.1385421.

[11] Anne-Mette Sauër, Cornelis Kalkman, and Diederik van Dijk. Postoperative cognitive decline. Journal of Anesthesia, 23(2):256–259, may 2009. ISSN 1438-8359. doi: 10.1007/s00540-009-0744-5. URL https://doi.org/10.1007/s00540-009-0744-5.

[12] Danielle Greaves, Peter J. Psaltis, Tyler J. Ross, Daniel Davis, Ashleigh E. Smith, Monique S. Boord, and Hannah A. D. Keage. Cognitive outcomes following coronary artery bypass grafting: A systematic review and meta-analysis of 91,829 patients. International Journal of Cardiology, 289:43–49, 2019. ISSN 0167-5273. doi: 10.1016/j.ijcard.2019.04.065. URL https://doi.org/10.1016/j.ijcard.2019.04.065.

[13] Lisbeth Evered, Brendan Silbert, David S. Knopman, David A. Scott, Steven T. DeKosky, Lars S. Rasmussen, Esther S. Oh, Greg Crosby, Miles Berger, and Roderic G. Eckenhoff. Recommendations for the nomenclature of cognitive change associated with anaesthesia and surgery—2018. British Journal of Anaesthesia, 121(5):1005–1012, 2018. doi: 10.1016/j.bja.2017.11.087.

[14] Lingyan Liang, Yueming Yuan, Yichen Wei, Bihan Yu, Wei Mai, Gaoxiong Duan, Xiucheng Nong, Chong Li, Jiahui Su, Lihua Zhao, Zhiguo Zhang, and Demao Deng. Recurrent and concurrent patterns of regional BOLD dynamics and functional connectivity dynamics in cognitive decline. Alzheimer’s Research & Therapy, 13(1):28, jan 2021. ISSN 1758-9193. doi: 10.1186/s13195-020-00764-6. URL https://doi.org/10.1186/s13195-020-00764-6.

[15] William W. Seeley, Vinod Menon, Alan F. Schatzberg, Jennifer Keller, Gary H. Glover, Heather Kenna, Allan L. Reiss, and Michael D. Greicius. Dissociable intrinsic connectivity networks for salience processing and executive control. Journal of Neuroscience, 27(9):2349–2356, 2007. ISSN 0270-6474. doi: 10.1523/JNEUROSCI.5587-06.2007. URL https://www.jneurosci.org/content/27/9/2349.

[16] Randy L. Buckner, Jessica R. Andrews-Hanna, and Daniel L. Schacter. The brain’s default network: Anatomy, function, and relevance to disease. In The year in cognitive neuroscience 2008, Annals of the New York Academy of Sciences, pages 1–38. Blackwell Publishing, Malden, 2008. ISBN 978-1-57331-726-9. ID: 2008-05643-001.

[17] Gustavo Deco, Viktor K. Jirsa, and Anthony R. McIntosh. Emerging concepts for the dynamical organization of resting-state activity in the brain. Nature Reviews Neuroscience, 12(1):43–56, 2011. doi: 10.1038/nrn2961.

[18] R. Matthew Hutchison, Thilo Womelsdorf, Evan A. Allen, Peter A. Bandettini, Vince D. Calhoun, Maurizio Corbetta, Stefania Della Penna, Jeff H. Duyn, Gary H. Glover, Javier Gonzalez-Castillo, Daniel A. Handwerker, Shella Keilholz, Vesa Kiviniemi, David A. Leopold, Francesco de Pasquale, Olaf Sporns, Markus Walter, and Catie Chang. Dynamic functional connectivity: promise, issues, and interpretations. NeuroImage, 80:360–378, 2013. doi: 10.1016/j.neuroimage.2013.05.079.

[19] Gopikrishna Deshpande, Stephan LaConte, George Andrew James, Scott Peltier, and Xiaoping Hu. Multivariate granger causality analysis of fmri data. Human Brain Mapping, 30(4):1361–1373, 2009. doi: 10.1002/hbm.20606. URL https://onlinelibrary.wiley.com/doi/abs/10.1002/hbm.20606.

[20] Daniel J. Lurie, Daniel Kessler, Danielle S. Bassett, Richard F. Betzel, Michael Breakspear, Shella Kheilholz, Aaron Kucyi, Raphaël Liégeois, Martin A. Lindquist, Anthony Randal McIntosh, Russell A. Poldrack, James M. Shine, William Hedley Thompson, Natalia Z. Bielczyk, Linda Douw, Dominik Kraft, Robyn L. Miller, Muthuraman Muthuraman, Lorenzo Pasquini, Adeel Razi, Diego Vidaurre, Hua Xie, and Vince D. Calhoun. Questions and controversies in the study of time-varying functional connectivity in resting fmri. Network Neuroscience, 4(1):30–69, 02 2020. ISSN 2472-1751. doi: 10.1162/netna00116. URL https://doi.org/10.1162/netn_a_00116.

[21] David Meunier, Renaud Lambiotte, and Edward T. Bullmore. Modular and hierarchically modular organization of brain networks. Frontiers in Neuroscience, Volume 4-2010, 2010. ISSN 1662-453X. doi: 10.3389/fnins.2010.00200. URL https://www.frontiersin.org/journals/neuroscience/articles/10.3389/fnins.2010.00200.

[22] Ed Bullmore and Olaf Sporns. The economy of brain network organization. Nature Reviews Neuroscience, 13(5):336–349, May 2012. ISSN 1471-0048. doi: 10.1038/nrn3214. URL https://doi.org/10.1038/nrn3214.

[23] Richard F. Betzel and Danielle S. Bassett. Multi-scale brain networks. NeuroImage, 160:73–83, 2017. ISSN 1053-8119. doi: 10.1016/j.neuroimage.2016.11.006. URL https://www.sciencedirect.com/science/article/pii/S1053811916306152. Functional Architecture of the Brain.

[24] Bei Rong, Huan Huang, Guoqing Gao, Limin Sun, Yuan Zhou, Ling Xiao, Huiling Wang, and Gaohua Wang. Widespread intra-and inter-network dysconnectivity among large-scale resting state networks in schizophrenia. Journal of Clinical Medicine, 12(9), 2023. ISSN 2077-0383. doi: 10.3390/jcm12093176. URL https://www.mdpi.com/2077-0383/12/9/3176.

[25] Erfan Nozari, Maxwell A. Bertolero, Jennifer Stiso, Lorenzo Caciagli, Eli J. Cornblath, Xiaosong He, Arun S. Mahadevan, George J. Pappas, and Dani S. Bassett. Macroscopic resting-state brain dynamics are best described by linear models. Nature Biomedical Engineering, 8(1):68–84, 2024. ISSN 2157-846X. doi: 10.1038/s41551-023-01117-y. URL https://doi.org/10.1038/s41551-023-01117-y.

[26] Lucia Colombo, Giuseppe Sartori, and Cristina Brivio. Stima del quoziente intellettivo tramite l’applicazione del tib (test breve di intelligenza). Giornale italiano di psicologia, Rivista trimestrale, (3/2002):613–638, 2002. ISSN 0390-5349. doi: 10.1421/1256. URL https://www.rivisteweb.it/doi/10.1421/1256.

[27] Christopher Randolph, Michael C. Tierney, Erich Mohr, and Thomas N. Chase. The repeatable battery for the assessment of neuropsychological status (RBANS): Preliminary clinical validity. Journal of Clinical and Experimental Neuropsychology, 20(3):310–319, 1998. ISSN 1380-3395. doi: 10.1076/jcen.20.3.310.823. URL https://doi.org/10.1076/jcen.20.3.310.823.

[28] Susan Whitfield-Gabrieli and Alfonso Nieto-Castanon. Conn: A functional connectivity toolbox for correlated and anticorrelated brain networks. Brain Connectivity, 2(3):125–141, 2012. doi: 10.1089/brain.2012.0073.

[29] Yashar Behzadi, Khaled Restom, Joy Liau, and Thomas T. Liu. A component based noise correction method (compcor) for bold and perfusion based fmri. NeuroImage, 37(1):90–101, 2007. ISSN 1053-8119. doi: 10.1016/j.neuroimage.2007.04.042. URL https://www.sciencedirect.com/science/article/pii/S1053811907003837.

[30] Michael N. Hallquist, Kai Hwang, and Beatriz Luna. The nuisance of nuisance regression: Spectral misspecification in a common approach to resting-state fmri preprocessing reintroduces noise and obscures functional connectivity. NeuroImage, 82:208–225, 2013. ISSN 1053-8119. doi: 10.1016/j.neuroimage.2013.05.116. URL https://www.sciencedirect.com/science/article/pii/S1053811913006265.

[31] Thomas M Cover. Elements of information theory. John Wiley & Sons, 1999.

[32] Tyrone E Duncan. On the calculation of mutual information. SIAM Journal on Applied Mathematics, 19(1):215–220, 1970.

[33] Lionel Barnett, Adam B. Barrett, and Anil K. Seth. Granger causality and transfer entropy are equivalent for gaussian variables. Physical Review Letters, 103(23):238701, 2009. doi: 10.1103/PhysRevLett.103.238701.

[34] Yuri Antonacci, Chiara Barà, Andrea Zaccaro, Francesca Ferri, Riccardo Pernice, and Luca Faes. Time-varying information measures: an adaptive estimation of information storage with application to brain-heart interactions. Frontiers in Network Physiology, 3:1242505, 2023. ISSN 2674-0109. doi: 10.3389/fnetp.2023.1242505. URL https://www.frontiersin.org/journals/network-physiology/articles/10.3389/fnetp.2023.1242505.

[35] Laura Sparacino, Yuri Antonacci, Chiara Barà, Angela Valenti, Alberto Porta, and Luca Faes. A method to assess granger causality, isolation and autonomy in the time and frequency domains: Theory and application to cerebrovascular variability. IEEE Transactions on Biomedical Engineering, 71(5):1454–1465, 2024. doi: 10.1109/TBME.2023.3340011.

[36] Alexander Kraskov, Harald Stögbauer, and Peter Grassberger. Estimating mutual information. Phys. Rev. E, 69:066138, Jun 2004. doi: 10.1103/PhysRevE.69.066138. URL https://link.aps.org/doi/10.1103/PhysRevE.69.066138.

[37] Valeria Rosalia Vergara, Chiara Barà, Andrea Zaccaro, Francesca Ferri, Fabrice Jurysta, Luca Faes, and Yuri Antonacci. Information-theoretic analysis of eeg wave amplitude and heart rate variability reveals the time scale-dependent nature of brain–heart interactions. IEEE Open Journal of Engineering in Medicine and Biology, 6:499–506, 2025. doi: 10.1109/OJEMB.2025.3590598.

[38] Wanting Xiong, Luca Faes, and Plamen Ch. Ivanov. Entropy measures, entropy estimators, and their performance in quantifying complex dynamics: Effects of artifacts, nonstationarity, and long-range correlations. Phys. Rev. E, 95:062114, Jun 2017. doi: 10.1103/PhysRevE.95.062114. URL https://link.aps.org/doi/10.1103/PhysRevE.95.062114.

[39] Laura Sparacino, Luca Faes, Gorana Mijatović, Giuseppe Parla, Vincenzina Lo Re, Roberto Miraglia, Jean de Ville de Goyet, and Gianvincenzo Sparacia. Statistical approaches to identify pairwise and high-order brain functional connectivity signatures on a single-subject basis. Life, 13(10):2075, 2023. ISSN 2075-1729. doi: 10.3390/life13102075. URL https://doi.org/10.3390/life13102075.

[40] Mikail Rubinov and Olaf Sporns. Complex network measures of brain connectivity: uses and interpretations. NeuroImage, 52(3):1059–1069, 2010. doi: 10.1016/j.neuroimage.2009.10.003.

[41] Jeffrey N. Browndyke, Miles Berger, Todd B. Harshbarger, Patrick J. Smith, William White, Tiffany L. Bisanar, John H. Alexander, Jeffrey G. Gaca, Kathleen Welsh-Bohmer, Mark F. Newman, and Joseph P. Mathew. Resting-state functional connectivity and cognition after major cardiac surgery in older adults without preoperative cognitive impairment: Preliminary findings. Journal of the American Geriatrics Society, 65(1):e6–e12, 2017. doi: 10.1111/jgs.14534. URL https://agsjournals.onlinelibrary.wiley.com/doi/abs/10.1111/jgs.14534.

[42] Emilio J. Sanz-Arigita, Menno M. Schoonheim, Jessica S. Damoiseaux, Serge A. R. B. Rombouts, Esther Maris, Frederik Barkhof, Philip Scheltens, and Cornelis J. Stam. Loss of ‘small-world’ networks in alzheimer’s disease: graph analysis of fmri resting-state functional connectivity. PLoS ONE, 5(11): e13788, 2010. doi: 10.1371/journal.pone.0013788.

[43] Betty M. Tijms, Alle Meije Wink, Willem de Haan, Wiesje M. van der Flier, Cornelis J. Stam, Philip Scheltens, and Frederik Barkhof. Alzheimer’s disease: connecting findings from graph theoretical studies of brain networks. Neurobiology of Aging, 34(8):2023–2036, 2013. doi: 10.1016/j.neurobiolaging.2013.02.020.

[44] Vinod Menon. Large-scale brain networks and psychopathology: a unifying triple network model. Trends in Cognitive Sciences, 15(10):483–506, 2011. ISSN 1364-6613. doi: 10.1016/j.tics.2011.08.003. URL https://www.sciencedirect.com/science/article/pii/S1364661311001719.

[45] Maurizio Corbetta and Gordon L. Shulman. Control of goal-directed and stimulus-driven attention in the brain. Nature Reviews Neuroscience, 3(3):201–215, mar 2002. ISSN 1471-0048. doi: 10.1038/nrn755. URL https://doi.org/10.1038/nrn755.

[46] Niccolò Terrando Lars I. Eriksson, Jae Kyu Ryu, Ting Yang, Claudia Monaco, Marc Feldmann, Malin Jonsson Fagerlund, Israel F. Charo, Katerina Akassoglou, and Mervyn Maze. Resolving postoperative neuroinflammation and cognitive decline. Annals of Neurology, 70(6):986–995, 2011. doi: 10.1002/ana.22664. URL https://onlinelibrary.wiley.com/doi/abs/10.1002/ana.22664.

[47] Sophie Achard and Ed Bullmore. Efficiency and cost of economical brain functional networks. PLoS Computational Biology, 3(2):e17, 2007. doi: 10.1371/journal.pcbi.0030017.

[48] Nicolas A. Crossley, Andrea Mechelli, James Scott, Francesco Carletti, Philip T. Fox, Philip McGuire, and Edward T. Bullmore. The hubs of the human connectome are generally implicated in the anatomy of brain disorders. Brain, 137(8):2382–2395, 2014. doi: 10.1093/brain/awu132.

[49] Cornelis Jan Stam. Hub overload and failure as a final common pathway in neurological brain network disorders. Network Neuroscience, 8(1):1–23, 04 2024. ISSN 2472-1751. doi: 10.1162/netna00339. URL https://doi.org/10.1162/netn_a_00339.

[50] Cornelis J. Stam and Elisabeth C. W. van Straaten. The organization of physiological brain networks. Clinical Neurophysiology, 123(6):1067–1087, 2012. doi: 10.1016/j.clinph.2012.01.011.

[51] Laura Sparacino, Yuri Antonacci, Gorana Mijatovic, and Luca Faes. Measuring hierarchically-organized interactions in dynamic networks through spectral entropy rates: Theory, estimation, and illustrative application to physiological networks. Neurocomputing, 630:129675, 2025. ISSN 0925-2312. doi: 10.1016/j.neucom.2025.129675. URL https://www.sciencedirect.com/science/article/pii/S0925231225003479.

[52] François Paugam, Basile Pinsard, Guillaume Lajoie, and Pierre Bellec. A benchmark of individual auto-regressive models in a massive fmri dataset. Imaging Neuroscience, 2:imag–2–00228, 07 2024. ISSN 2837-6056. doi: 10.1162/imaga00228. URL https://doi.org/10.1162/imag_a_00228.

