## Supplementary for "Information-Theoretic Functional Connectivity Characterizes Multiscale Network Reorganization in Postoperative Cognitive Decline"

| Resting-state network | Brain region | MNI coordinates in mm (X,Y,Z) |
|---|---|---|
| Default Mode | Medial prefrontal cortex | 1 55 -3 |
| Default Mode | Left lateral parietal | -39 -77 33 |
| Default Mode | Right lateral parietal | 47 -67 29 |
| Default Mode | Posterior cingulate cortex | 1 -61 38 |
| Sensori Motor | Left lateral sensorimotor | -55 -12 29 |
| Sensori Motor | Right lateral Sensorimotor | 56 -10 29 |
| Sensori Motor | Superior sensorimotor | 0 -31 67 |
| Visual | Medial visual | 2 -79 12 |
| Visual | Occipital visual | 0 -93 -4 |
| Visual | Left lateral visual | -37 -79 10 |
| Visual | Right lateral visual | 38 -72 13 |
| Salience | Anterior cingulate cortex | 0 22 35 |
| Salience | Left anterior insula | -44 13 1 |
| Salience | Right anterior insula | 47 14 0 |
| Salience | Left rostral prefrontal cortex | -32 45 27 |
| Salience | Right rostral prefrontal cortex | 32 46 27 |
| Salience | Left supramarginal gyrus | -60 -39 31 |
| Salience | Right supramarginal gyrus | 62 -35 32 |
| Dorsal Attention | Left frontal eye field | -27 -9 64 |
| Dorsal Attention | Right frontal eye field | 30 -6 64 |
| Dorsal Attention | Left intraparietal sulcus | -39 43 52 |
| Dorsal Attention | Right intraparietal sulcus | 39 -42 54 |
| Fronto Parietal | Left lateral prefrontal cortex | -43 33 28 |
| Fronto Parietal | Left posterior parietal cortex | -46 -58 49 |
| Fronto Parietal | Right lateral prefrontal cortex | 41 38 30 |
| Fronto Parietal | Right posterior parietal cortex | 52 -52 45 |
| Language | Left inferior frontal gyrus | -51 26 2 |
| Language | Right inferior frontal gyrus | 54 28 1 |
| Language | Left posterior superior temporal gyrus | -57 -47 15 |
| Language | Right posterior superior temporal gyrus | 59 -42 13 |
| Cerebellar | Anterior cerebellar | 0 -63 -30 |
| Cerebellar | Posterior cerebellar | 0 -79 -32 |

Table S1: Primary resting-state networks used as regions of interest (ROIs), with corresponding brain regions and MNI coordinates (X, Y, Z in mm).

| | m = 1 | | m = 5 | |
|---|---|---|---|---|
| | MIR linear | MIR knn | MIR linear | MIR knn |
| Global FC | 0.187 | 0.421 | 0.192 | 0.208 |
| Intra-RSN | | | | |
| DM | 0.821 | 0.679 | 0.592 | 0.275 |
| SM | 0.968 | 0.123 | 0.132 | 0.922 |
| VS | 0.649 | 0.721 | 0.521 | 0.168 |
| SAL | 0.159 | 0.581 | 0.155 | 0.715 |
| DA | 0.312 | 0.385 | 0.413 | 0.076 |
| FP | 0.493 | 0.919 | 0.978 | 0.738 |
| L | 0.391 | 0.059 | 0.360 | 0.366 |
| CB | 0.522 | 0.418 | 0.937 | 0.488 |
| Inter-RSN | | | | |
| DM | 0.276 | 0.815 | 0.279 | 0.087 |
| SM | 0.542 | 0.147 | 0.341 | 0.716 |
| VS | 0.064 | 0.357 | 0.153 | 0.354 |
| SAL | 0.358 | 0.558 | 0.511 | 0.829 |
| DA | 0.342 | 0.316 | 0.381 | 0.132 |
| FP | 0.394 | 0.879 | 0.357 | 0.395 |
| L | 0.555 | 0.937 | 0.463 | 0.088 |
| CB | 0.066 | 0.827 | 0.125 | 0.123 |

Table S2: Results of permutation testing (p-values) comparing longitudinal functional connectivity changes ($\Delta FC = \mathrm{FU} - \mathrm{BL}$) between POCD and NO-POCD groups for Mutual Information Rate (MIR) linear and knn with $m = 1$ and $m = 5$ at global, intra-RSN, and inter-RSN scales.

| Group | Phase | MI linear | MI knn | MIR linear | MIR knn |
|---|---|---|---|---|---|
| **Overall** | Baseline | 126.21 ± 37.38 | 60.79 ± 41.24 | 241.50 ± 49.74 | 57.43 ± 44.62 |
| | Follow-Up | 140.71 ± 35.99 | 55.86 ± 17.81 | 227.86 ± 23.82 | 38.86 ± 7.93 |
| **NO POCD** | Baseline | 113.57 ± 12.26 | 47.29 ± 11.06 | 231.57 ± 44.78 | 47.57 ± 10.41 |
| | Follow-Up | 149.71 ± 48.14 | 57.86 ± 21.75 | 238.29 ± 17.60 | 42.29 ± 9.05 |
| **POCD** | Baseline | 138.86 ± 50.04 | 74.29 ± 56.02 | 251.43 ± 55.90 | 67.29 ± 63.08 |
| | Follow-Up | 131.71 ± 17.31 | 53.86 ± 14.30 | 217.43 ± 25.80 | 35.43 ± 5.19 |

Table S3: Mean (± standard deviation) number of ROI pairs with significant functional connectivity after surrogate analysis, estimated via linear and knn-based Mutual Information (MI) and Mutual Information Rate (MIR; $m = 3$). Results are reported for the overall cohort ($N = 14$), NO POCD ($N = 7$) and POCD ($N = 7$) groups at Baseline and Follow-Up.

| Variable | MI linear | MI knn | MIR linear | MIR knn |
|---|---|---|---|---|
| Δ Degree | **0.55 (0.046)** | 0.34 (0.226) | 0.30 (0.310) | 0.22 (0.494) |
| Δ Strength | 0.43 (0.106) | 0.30 (0.286) | 0.20 (0.498) | 0.28 (0.376) |
| Δ Density | **0.56 (0.030)** | 0.33 (0.260) | 0.30 (0.310) | 0.23 (0.476) |
| Δ Betweenness Centrality | -0.24 (0.408) | -0.02 (0.907) | -0.48 (0.076) | **-0.65 (0.012)** |
| Δ Global Efficiency | **0.66 (0.012)** | 0.20 (0.501) | 0.05 (0.833) | 0.12 (0.679) |
| Δ Local Efficiency | 0.34 (0.234) | 0.30 (0.324) | 0.09 (0.737) | 0.23 (0.452) |
| Δ Modularity | **-0.72 (0.006)** | -0.51 (0.066) | -0.49 (0.082) | -0.43 (0.126) |
| Δ Small-World Index | **-0.62 (0.014)** | -0.57 (0.051) | -0.29 (0.332) | **-0.58 (0.030)** |

Table S4: Spearman's correlation coefficients ($\rho$) and corresponding $p$-values (reported in parentheses) for the associations between RBANS scores and longitudinal changes ($\Delta = \mathrm{FU} - \mathrm{BL}$) in graph-theoretical metrics. Metrics are derived from functional connectivity based on Mutual Information (MI) and Mutual Information Rate (MIR; $m = 3$), using linear and knn-based estimators. Significant associations ($p < 0.05$) are highlighted in bold.

| | m = 1 | | m = 5 | |
|---|---|---|---|---|
| **Variable** | **MIR linear** | **MIR knn** | **MIR linear** | **MIR knn** |
| Δ Significant Edges | 0.45 (0.154) | 0.22 (0.462) | 0.45 (0.154) | 0.12 (0.679) |
| Δ Degree | 0.45 (0.154) | 0.22 (0.462) | 0.45 (0.154) | 0.12 (0.679) |
| Δ Density | 0.45 (0.154) | 0.22 (0.462) | 0.45 (0.154) | 0.12 (0.679) |
| Δ Strength | 0.36 (0.204) | 0.10 (0.723) | 0.25 (0.368) | 0.14 (0.597) |
| Δ Betweenness Centrality | -0.07 (0.819) | 0.34 (0.266) | 0.38 (0.174) | -0.01 (0.975) |
| Δ Global Efficiency | 0.19 (0.500) | -0.02 (0.951) | 0.06 (0.827) | 0.14 (0.643) |
| Δ Local Efficiency | 0.04 (0.863) | -0.04 (0.903) | -0.02 (0.899) | 0.16 (0.575) |
| Δ Modularity | -0.50 (0.086) | -0.14 (0.595) | **-0.55 (0.024)** | 0.15 (0.663) |
| Δ Small-World Index | 0.22 (0.446) | -0.08 (0.809) | -0.32 (0.238) | -0.11 (0.693) |

Table S5: Spearman's correlation coefficients ($\rho$) and corresponding $p$-values (reported in parentheses) for the associations between RBANS scores and longitudinal changes ($\Delta = \mathrm{FU} - \mathrm{BL}$) in graph-theoretical metrics. Metrics are derived from functional connectivity based on Mutual Information Rate (MIR), using linear and knn-based estimators with $m = 1$ and $m = 5$. Significant associations ($p < 0.05$) are highlighted in bold.

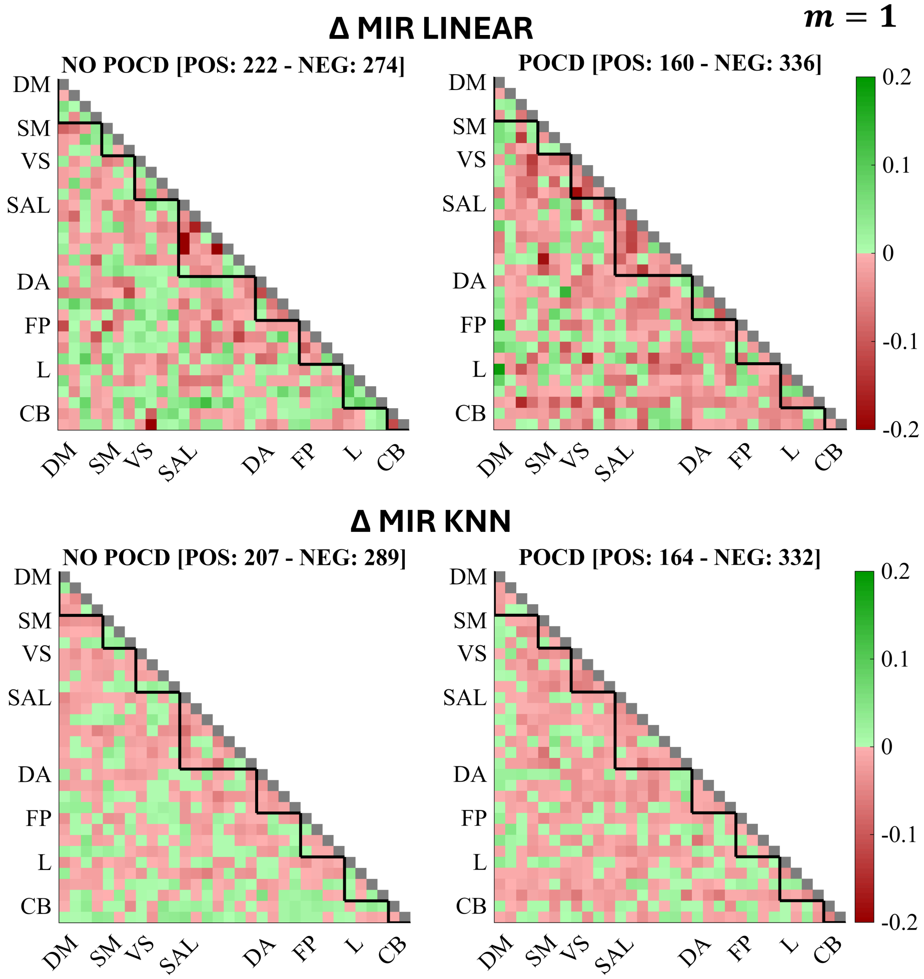


Figure S1: Median Δ FC matrices for the two groups NO POCD (left column) and POCD (right column) estimated as linear (top row) and knn-based model-free (bottom row) MIR with $m = 1$. Edges with a positive difference are reported in green, while edges with negative changes in red. Number of positive (POS) and negative (NEG) edges are reported in the figure. Black triangles show intra-RSN FC portion.

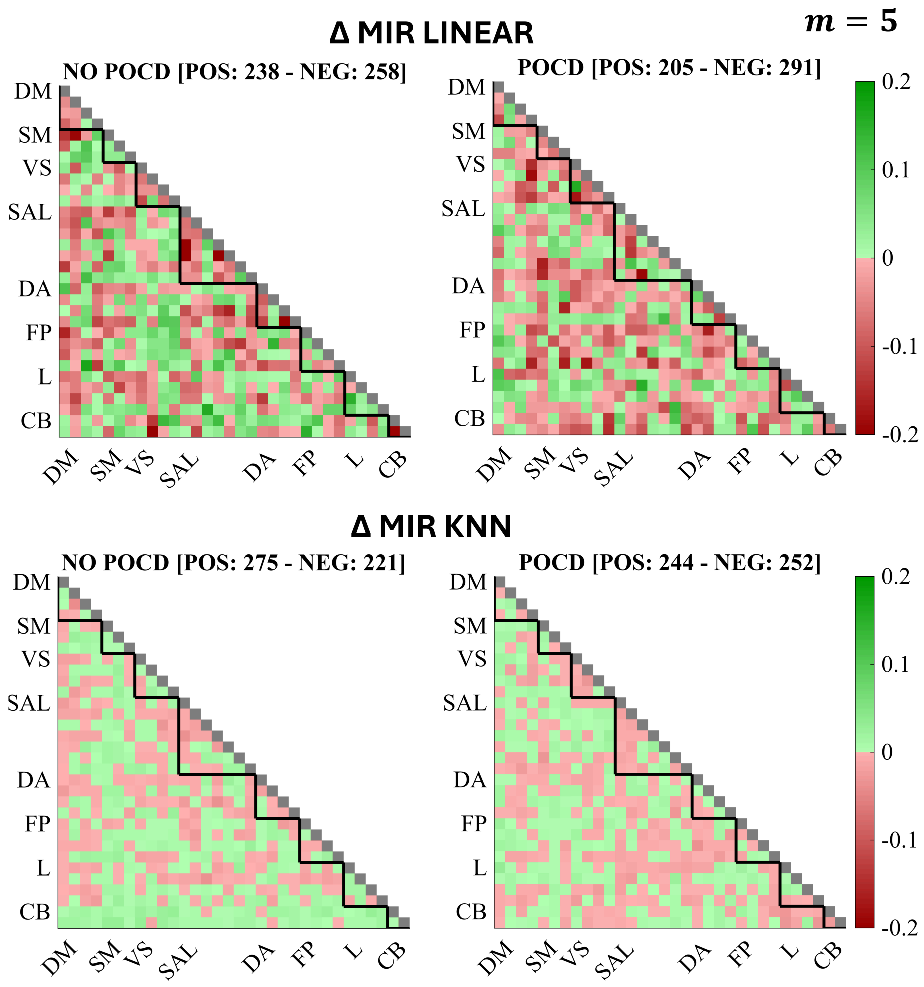


Figure S2: Median Δ FC matrices for the two groups NO POCD (left column) and POCD (right column) estimated as linear (top row) and knn-based model-free (bottom row) MIR with $m = 5$. Edges with a positive difference are reported in green, while edges with negative changes in red. Number of positive (POS) and negative (NEG) edges are reported in the figure. Black triangles show intra-RSN FC portion.
